# Modulation of adenine phosphoribosyltransferase-mediated salvage to promote diabetic wound healing

**DOI:** 10.1101/2020.04.08.032128

**Authors:** Guang-Huar Young, Jiun-Tsai Lin, Yi-Fang Cheng, Chia-Fang Ho, Qian-Yu Kuok, Ru-Chun Hsu, Wan-Rou Liao, Chin-Chen Chen, Han-Min Chen

## Abstract

Adenine phosphoribosyltransferase (APRT) is the key enzyme in purine salvage by the incorporation of adenine and phosphoribosyl pyrophosphate to provide adenylate nucleotide. The up-regulated APRT found in wound skin correlated with the demands of repair in diabetic mice. Administration of adenine on the wound of diabetic mice exhibited elevated ATP levels in organismic skin and accelerated wound healing. In vitro studies showed that APRT utilized adenine to rescue cellular ATP levels and proliferation against hydrogen peroxide-induced oxidative damage. LC-MS/MS-based analysis of total adenylate nucleotides in NIH-3T3 fibroblast showed that adenine addition enlarged the cellular adenylate pool, reduced the adenylate energy charge, and provided more AMP for the generation of ATP in further. These data indicated the role of APRT during diabetic wound healing by regulating the nucleotide pool after injury and demonstrated the improvement by topical adenine, which highlights its value as a promising agent in therapeutic intervention. Our study provided an explanation for the up- regulation of APRT in tissue repair and adenine supplement resulted in an enlargement of the adenylate pool for ATP generation.

## Introduction

The purine salvage pathway appears primarily in extrahepatic tissues (Bhagavan & Ha, 2011) and normally activates after wounding (Murray, 1971). The re-utilization of purine base requires less energy than *de novo* synthesis, accounting for 90% of daily purine nucleotide biosynthesis. Adenine phosphoribosyl transferase (APRT), which is the key enzyme in the salvage pathway for purine biosynthesis and is widely distributed throughout various organs of mammals (Murray, 1971). APRT catalyzes the synthesis of adenosine monophosphate (AMP) from adenine and the ribose derivative 5’-phosphoribosyl-1-pyrophosphate (Thimm, Schiedel et al., 2015). In cutaneous wound healing, both APRT and its anabolite AMP increase in the wounding site (Gassmann, Stanzel et al., 1999, Rossomando & Bertolami, 1983). Hence, the up-regulation of APRT might be for the supplement of adenylate nucleotides, which serve as energy donors for wound repair. However, the role of APRT in purine metabolism on the chronic wound, such as diabetic wound, remains poorly understood.

Tissue injury elevates the cellular energy demand and triggers quiescent cells to re-enter the cell cycle (Im & Hoopes, 1970a, Vande Berg & Robson, 2003). Physiologically, wound healing requires considerable events for cooperation and involves numerous cells, growth factors, cytokines, and enzymes (Falanga, 2005). These processes require extra energy expenditure, mainly in the form of ATP (Im & Hoopes, 1970a). In chronic wounds, tissue hypoxia results in energy supply deletion and oxidative damage, which is the cause of the chronic non-healing wounds (Cano Sanchez, Lancel et al., 2018). Therefore, lots of wound managements show that antioxidant strategies or direct energy supply by exogenous ATP-liposome accelerates diabetic wound healing through elevating the cellular ATP levels (Howard, Sarojini et al., 2014). However, the only one drug approved by FDA for the treatment of diabetic wound is associated with the risk of cancer mortality and expensive concerns (Papanas & Maltezos, 2010). This highlighted the medical need for diabetic wound drugs to improve the quality of life with reduced treatment costs.

AMP-activated protein kinase (AMPK) modulating energy metabolism may potentially target the strong association between cellular energy supply and various diseases, such as diabetes, cancer, and cardiovascular disease (Hardie, 2015, Lin, Chen et al., 2014). Exogenous adenine has shown the potential to modulate the cellular activity of AMPK via APRT anabolism (Leu, Chiang et al., 2017, Young, Lin et al., 2015). In normal condition, the intracellular concentration of the adenine base is low when compared with other bases and nucleosides (Traut, 1994). We know that the only cellular source of free adenine base is derived from the by-product of the polyamine biosynthesis pathway (Kamatani & Carson, 1981) and is predominantly metabolized by APRT to yield AMP as the product (Murray, 1971). The application of adenine had been utilized in preserving tissue and blood transfusion products by cellular ATP replenishment (Foker, 1988, Frenguelli, 2017). Reasonably, the improvement of energy production by adenine supplement is beneficial for cells to overcome the energy threshold (Wang, Wan et al., 2010).

This study aims to investigate the role of APRT during skin wound repair. Tissue repair requires energy; thus, we provided a method to replenish cellular ATP levels to accelerate diabetic wound healing.

## Results

### Skin wound healing stimulated the expression of APRT in mice

To clarify the role of APRT on wound repair duration, we first used a full-thickness excisional wound model in both wild-type mice (C57BL/6) and diabetic (db/db) mice. The wound areas were recorded using a digital camera daily, as shown in Figure 1A. The overall closure time in wild-type mice was more rapid than that in diabetic mice. Most of the wounds in wild-type mice were closed after 13 days, whereas as those in diabetic mice would completely close nearly double time with the same area size. The average of wound closure time in diabetic mice was 21.9 ± 1.3 days. Western blot analysis of wild-type mice shown in Supplemental Figure 1 revealed that the APRT protein levels in a wounded skin exhibited a maximum increase on day 3 after wounding compare with those in an unwounded skin as control (1.4 ± 0.2 vs. 0.5 ± 0.1, P < 0.05). However, on day10, the APRT protein levels in a wounded skin after wounding from wild-type mice, the levels of APRT protein in the wound region subsequently declined. Conversely, in diabetic mice, the APRT protein levels in a wounded skin sequentially increase after wounding compared with those control skin, and the expression of APRT protein still up- regulated in the closed diabetic wounds compared to control skin (day 20: 0.9 ± 0.1 vs. 0.3 ± 0.1, P < 0.05), as shown in Figure 1B and 1C. Immunohistological staining of a diabetic mouse skin showed that the APRT predominantly elevated in the epidermal layer of the wounded skin, especially in dermal fibroblasts and keratinocytes (Figure 1D).

**Fig. 1.**
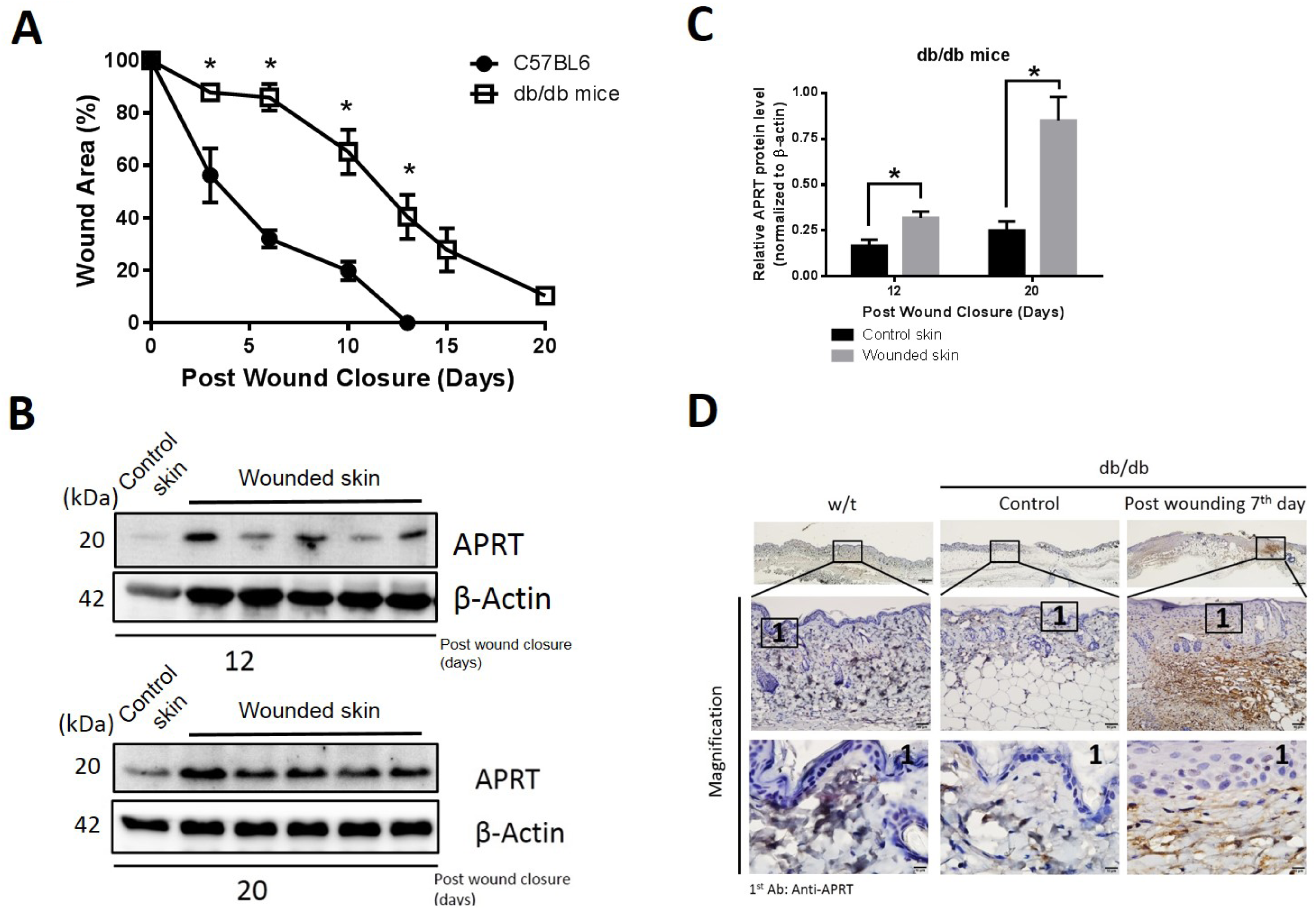
Expression of adenine phosphoribosyltransferase (APRT) during cutaneous wound repair in diabetic mice. Wild-type mice and diabetic mice were wounded as described in the materials and methods section. (A) Change of wound area on the duration of wound healing. (B) Representative Western blots for APRT in the diabetic mice skin at an indicated time point; β-actin was used as loading control. (C) Quantitation of the intensity of Western blots for APRT. Data were expressed as mean ± SEM, N ≥ 5. *, P < 0.05; N.S., not-significant. (D) APRT expression in the epidermal skin of wild-type and diabetic mice. Histological sections were stained by immunohistochemistry with anti-APRT antibodies and were counterstained with hematoxylin. Scale bars, 500 μm.

### Adenine administration accelerated diabetic wound healing

Given that APRT elevation in a wounded skin may reflect the nucleotide demand for wound repair, we speculated that exogenous adenine as an APRT substrate in the severe wound could benefit for tissue repair, particularly in a diabetic wound. To minimize wound contraction, we employed a silicone-splinted excisional wound model to calculate the residual wound area and captured representative photographs daily after wounding (Figure 2A). Prior to these experiments, several adenine concentrations were pre-examined, and daily administration of 0.05% adenine on a topical wound revealed the best efficiency for diabetic wound healing (data not shown). Figure 2B illustrates a significant decrease in terms of average wound area in adenine-treated wounds in comparison with that in the vehicle groups from day 6 after wounding (*, P < 0.05). The average unhealed wound area was 48.2% ± 5.3% of the initial wound area in the adenine-treated group, whereas 65.2% ± 5.5% was obtained in the vehicle group on day 6 after wounding. On average, it took 18.3 ± 1.0 days for adenine-treated wounds to close completely in contrast with 21.9 ± 1.3 days for vehicle groups (Figure 2C).

**Fig. 2.**
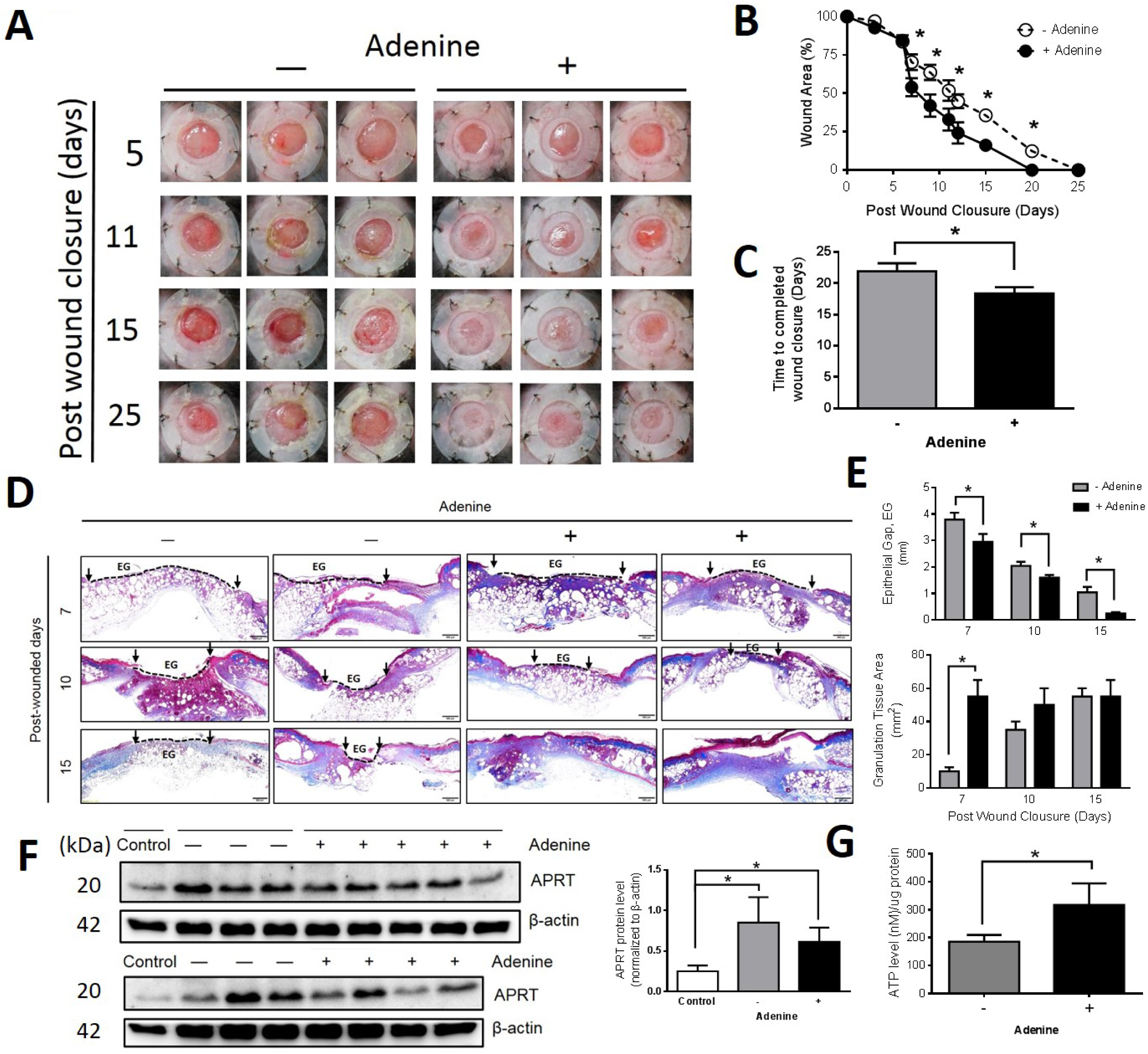
Diabetic wound healing following topical application of adenine. (A) Representative images of wounds in diabetic mice at different times after treatment. (B) Change of wound area on the duration of wound healing. (C) Comparison of wound healing completion time between the control and adenine group. (D) Cross sections of incised wounds were subjected to Masson’s trichrome staining. (E) Measurements of epithelial gaps and granulation tissue area in the wound. Data were expressed as mean ± SEM. *P < 0.05 vs. vehicle, N = 5. (F) Representative Western blots and quantification for APRT in the diabetic mice skin at day 20. β-Actin was used as loading control. (G) Relative ATP levels in diabetic wounds at day 20 post-injury. Data were expressed as mean ± SEM. *, P < 0.05; **, P < 0.01 vs. vehicle, N ≥ 5.

Re-epithelialization, granulation and collagen deposition are early markers of wound healing (Pastar, Stojadinovic et al., 2014). In Figure 2D, the histologic assessment of diabetic wounds confirmed that adenine treatment promoted those markers by Masson’s trichrome staining. The area size of adenine-treated wounds significantly decreased, as reflected by reduced epithelial gaps at day 7, 10, and 15 after wounding (Figure 2E, up-panel). By quantitative measurement, adenine treatment significantly promoted the formation of granulation tissue (Figure 2E, bottom panel), which was significantly larger and thicker than that of the vehicle.

The APRT protein levels on the diabetic wound skin at day 20 were quantified by Western blot analysis, and the ATP levels in wounded tissues were then measured. In Figure 2F, the APRT remained elevated compared with the unwounded skin (control skin). Interestingly, the elevation of APRT in the wounded skin from the adenine group was less than the vehicle group. Figure 2B and 2F showed that the APRT level correlated inversely with the recovery of wound healing in diabetic mice. Furthermore, the determination of ATP levels in the wounded skin showed that the adenine group had elevated ATP levels by 1.3 ± 0.1-fold in the tissue compared with the vehicle group (Figure 2G).

### Effects of exogenous adenine on cellular adenylate nucleotide pool

Considering that the fibroblast is the major repair cell in wounded skin (Andreea, Marieta et al., 2008), we used mouse NIH-3T3 fibroblasts as a model to quantify the dynamic conversion among adenylate nucleotides (AMP, ADP, and ATP) by HPLC-ESI-MS/MS after adenine treatment.

NIH-3T3 fibroblasts exposed to serial concentrations of adenine resulted in a substantial increase of adenylate nucleotides in a time/dose-dependent manner (Table 1). Importantly, the cellular AMP, not the ADP and ATP, was first elevated after 10 min of adenine incubation (*, P < 0.05) compared with that in control, and the elevation continued until the end of the study for 180 min. Meanwhile, the delayed elevations of ADP (&, P < 0.05) and ATP ($, P < 0.05) compared with those in the control group were predominantly present in 30 and 60 min, respectively. Thus, the first product of adenylate nucleotide after adenine addition is AMP.

**Table 1.** Time-course of the adenylate nucleotides response to adenine supplementation in NIH-3T3 fibroblasts. Absolute concentrations of adenylate nucleotides exposed to different amounts of isotope-labeled adenine as monitored by MRM methods; N=4. Data presented as mean±SD.

|  |  | Adenine (μM) |  |  |  |  |  |
| --- | --- | --- | --- | --- | --- | --- | --- |
|  |  | 0 | 5 | 10 | 50 | 100 | 500 |
| AMP (μM) | 10 min | 0.23±0.04 | 0.36±0.05* | 0.45±0.05* | 0.44±0.02* | 0.39±0.03* | 0.45±0.05* |
|  | 30 min | 0.36±0.06 | 0.67±0.13* | 0.66±0.13* | 0.72±0.11* | 0.7±0.08* | 0.83±0.16* |
|  | 60 min | 0.21±0.05 | 0.45±0.08* | 0.63±0.13* | 0.83±0.2* | 0.74±0.15* | 0.85±0.25* |
|  | 180 min | 0.25±0.05 | 0.41±0.06* | 0.58±0.23* | 0.66±0.18* | 0.77±0.05* | 0.98±0.12* |
| ADP (μM) | 10 min | 0.21±0.03 | 0.3±0.07 | 0.3±0.05 | 0.29±0.05 | 0.29±0.04 | 0.35±0.09 |
|  | 30 min | 0.25±0.04 | 0.43±0.1<sup>&</sup> | 0.47±0.12<sup>&</sup> | 0.47±0.08<sup>&</sup> | 0.61±0.08<sup>&</sup> | 0.66±0.15<sup>&</sup> |
|  | 60 min | 0.33±0.07 | 0.59±0.13<sup>&</sup> | 0.7±0.12<sup>&</sup> | 0.85±0.12<sup>&</sup> | 0.79±0.14<sup>&</sup> | 0.95±0.2<sup>&</sup> |
|  | 180 min | 0.41±0.05 | 0.62±0.06<sup>&</sup> | 0.72±0.18<sup>&</sup> | 0.72±0.12<sup>&</sup> | 0.83±0.06<sup>&</sup> | 0.97±0.09<sup>&</sup> |
| ATP (μM) | 10 min | 1.04±0.04 | 1.18±0.15 | 1.21±0.05 | 0.97±0.09 | 1.1±0.1 | 1.1±0.1 |
| | 30 min | 0.77±0.15 | 1.03±0.08<sup>\$</sup> | 0.89±0.11<sup>\$</sup> | 1.3±0.18<sup>\$</sup> | 1.62±0.28<sup>\$</sup> | 1.64±0.23<sup>\$</sup> |
| | 60 min | 1.1±0.27 | 1.42±0.12<sup>\$</sup> | 1.76±0.22<sup>\$</sup> | 2.04±0.42<sup>\$</sup> | 1.96±0.21<sup>\$</sup> | 2.17±0.31<sup>\$</sup> |
| | 180 min | 1.02±0.09 | 1.5±0.19<sup>\$</sup> | 1.39±0.1<sup>\$</sup> | 1.38±0.11<sup>\$</sup> | 1.5±0.21<sup>\$</sup> | 1.8±0.2<sup>\$</sup> |

\*, P<0.05; Significantly different from 0 uM adenine control in each time group.

Figure 4A depicts the timely changes of the total adenine nucleotide (AMP + ADP + ATP) levels in NIH-3T3 fibroblasts exposed to serial adenine concentrations. The total adenylate nucleotide from the control group increased progressively in a time/dose-dependent manner. After exposure for 180 min, despite a minor increase in ADP levels, the total adenine nucleotide slightly decrease than that exposure for 60 min.

**Fig. 3.**
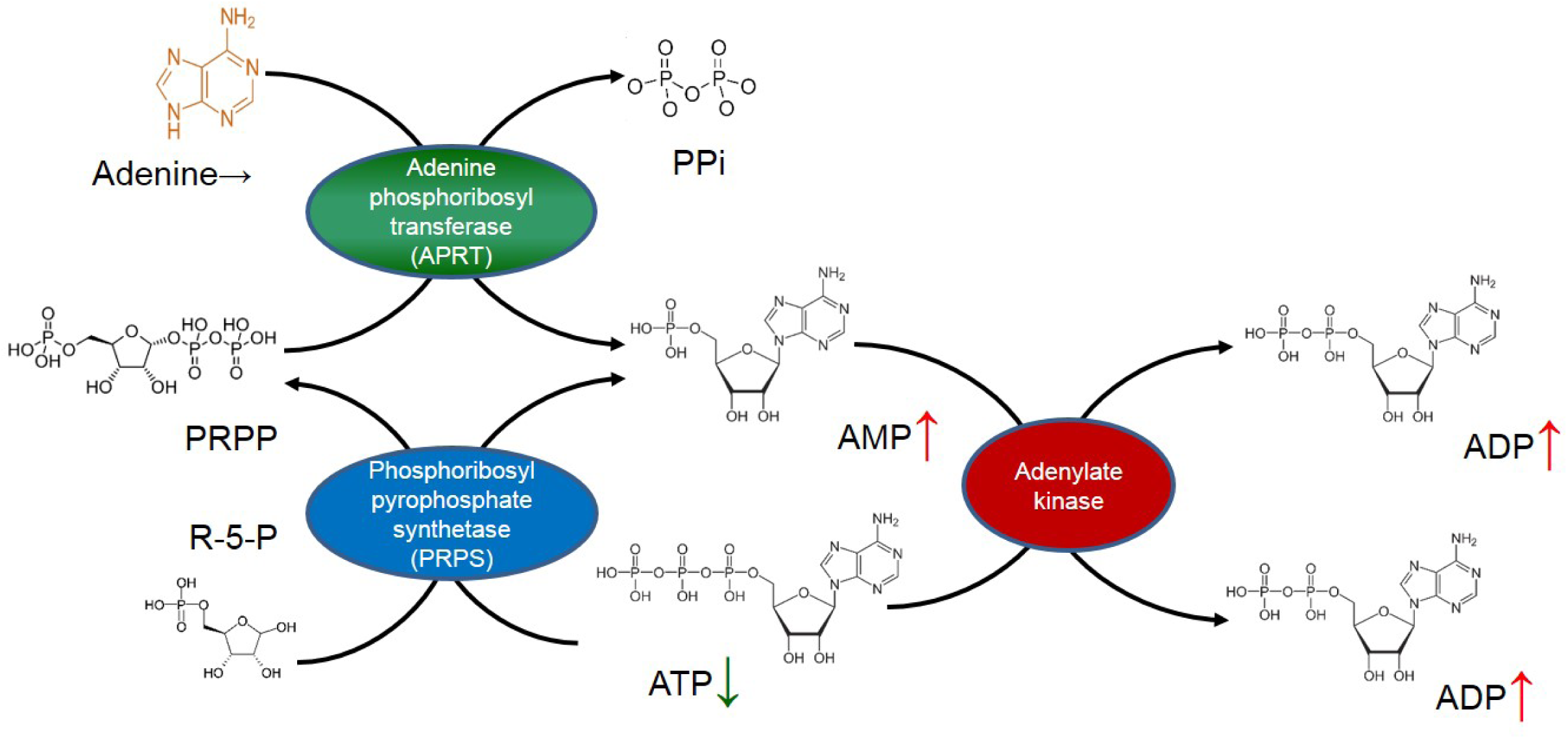
Schema illustrating the APRT-mediated salvage pathways for adenylate nucleotide conversion.

**Fig. 4.**
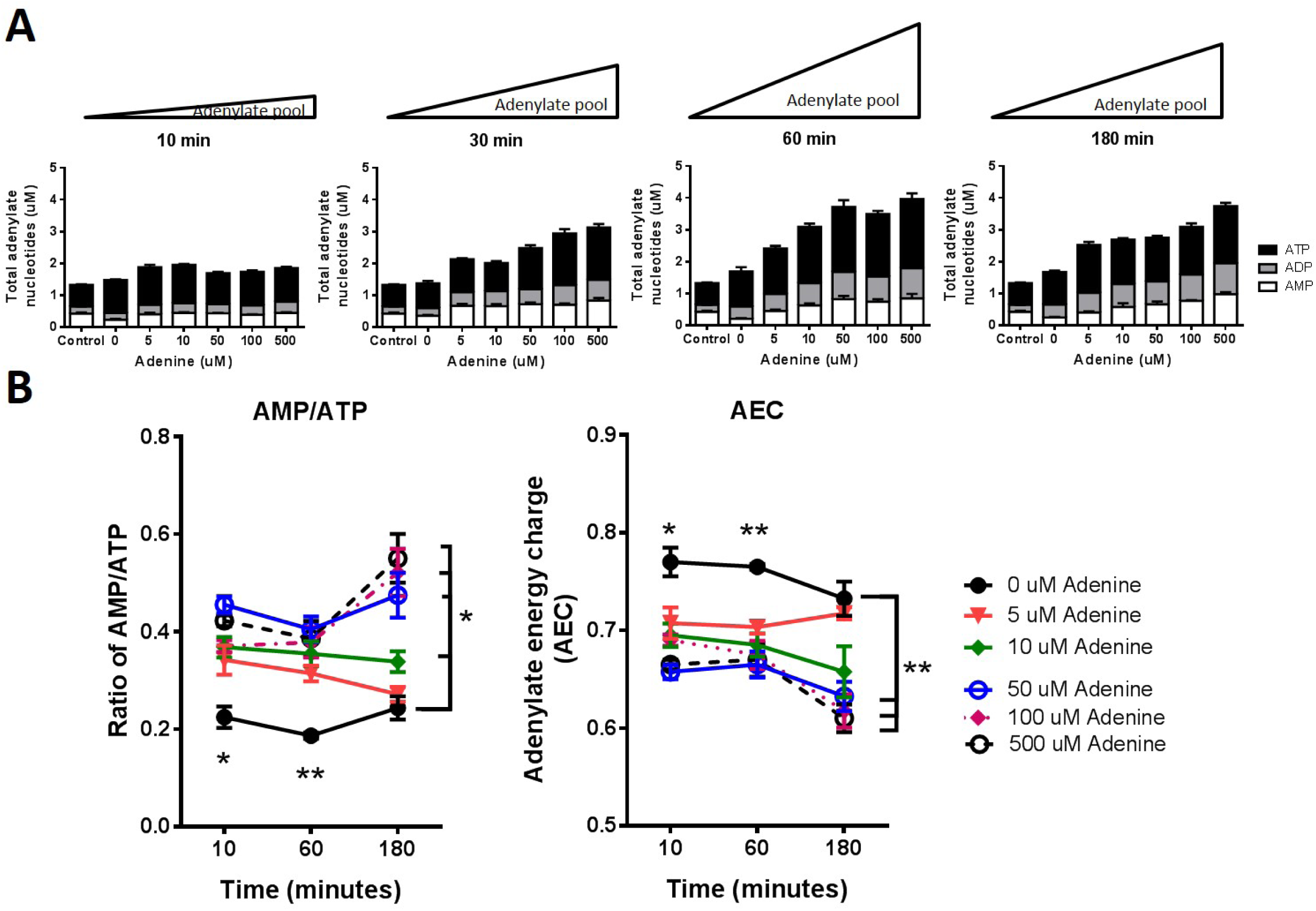
Adenine supplement enlarged the cellular adenylate pool in NIH-3T3 fibroblasts. Time course of the (A) total adenine nucleotide concentrations, (B) AMP / ATP ratio, and (C) adenylate energy charge response to adenine supplementation. Absolute concentrations of adenylate nucleotides exposed to different amounts of isotope-labeled adenine as monitored by MRM methods; N = 4. Data were presented as mean ± SEM.

To explore the response of the cellular energy state, we calculated the adenylate energy charge (AEC) values at each point. As defined by Atkinson and Walton (Atkinson, 1968), AEC represents the cellular energy status by the calculation of ([ATP] + 0.5[ADP])/([ATP] + [ADP] + [AMP]). *In vitro* studies showed that the AEC values stabilize in the range of 0.75-0.95 in normal cells, as predicted from kinetic studies. Values close to 1 indicate a health status, whereas values below 0.5 indicate the lack of energy in cells. Figure 4B shows that the AEC level values dropped from 0.77 ± 0.02 for the control cells to 0.71 ± 0.02 for the cells treated with low- dose adenine at 5 μmol after 10 min of treatment (P < 0.05); this phenomenon could be reversed to the control cellular level after 180 min. However, after exposing the cells to high-dose adenine greater or equal to 50 μmol for 180 min, the dramatic decline of the AEC level would be irreversible.

Despite the elevation of total adenylate nucleotides after adenine treatment, the increasing amplitude among AMP, ADP, and ATP compared with that in the control group at each time point was quite different. The AMP / ATP ratio is more functionally important than the absolute concentration of ATP, for instance, the regulation of AMPK activity (Hardie, 2004). Figure 4C illustrates that the AMP/ATP ratio increased immediately after adenine exposure for 10 min. Among the different doses of adenine groups, the low-dose adenine groups at 5 and 10 μmol gradually decreased back to the control levels following exposure for 180 min, but the high- dose adenine groups at 50, 100, and 500 μmol remained elevated continually. Interestingly, the AMP / ATP ratio showed a complementary distribution with the AEC value during the measurement.

### APRT mediated the cellular ATP levels to prevent H2O2 damage

The pathology of diabetic wound repair results from the overproduction of reactive oxygen species (ROS), leading to mitochondrial dysfunction and delaying the healing phase (Nouvong, Ambrus et al., 2016). To elucidate the role of APRT under the diabetic environment, we selected the haploid leukemia cell line HAP1 (Carette, Raaben et al., 2011) because it was available as an APRT-knockout cell line. Figure 5A shows that adenine elevated the cellular ATP levels of HAP1 cells in dose dependently. The expression of APRT protein was confirmed by Western blot, and Figure 5B reveals that APRT protein was absent in koAPRT HAP1 cells. Moreover, ATP analysis demonstrated that exogenous adenine-induced ATP elevation was absent in koAPRT HAP1 cells (Figure 5C). These data match the illustrating schema shown in Figure 3 in which adenine elevated cellular ATP is mediated by APRT. Hydrogen peroxide allowed the oxidative stress to mimic the diabetic condition, and the administration of 500 μmol adenine significantly reduced the hydrogen peroxide-induced decline of cellular ATP in wild-type cells; however, the effect of adenine was absent in koAPRT HAP1 cell lines. Therefore, APRT mediated exogenous adenine-induced ATP elevation against oxidative stress by introducing hydrogen peroxide.

**Fig. 5.**
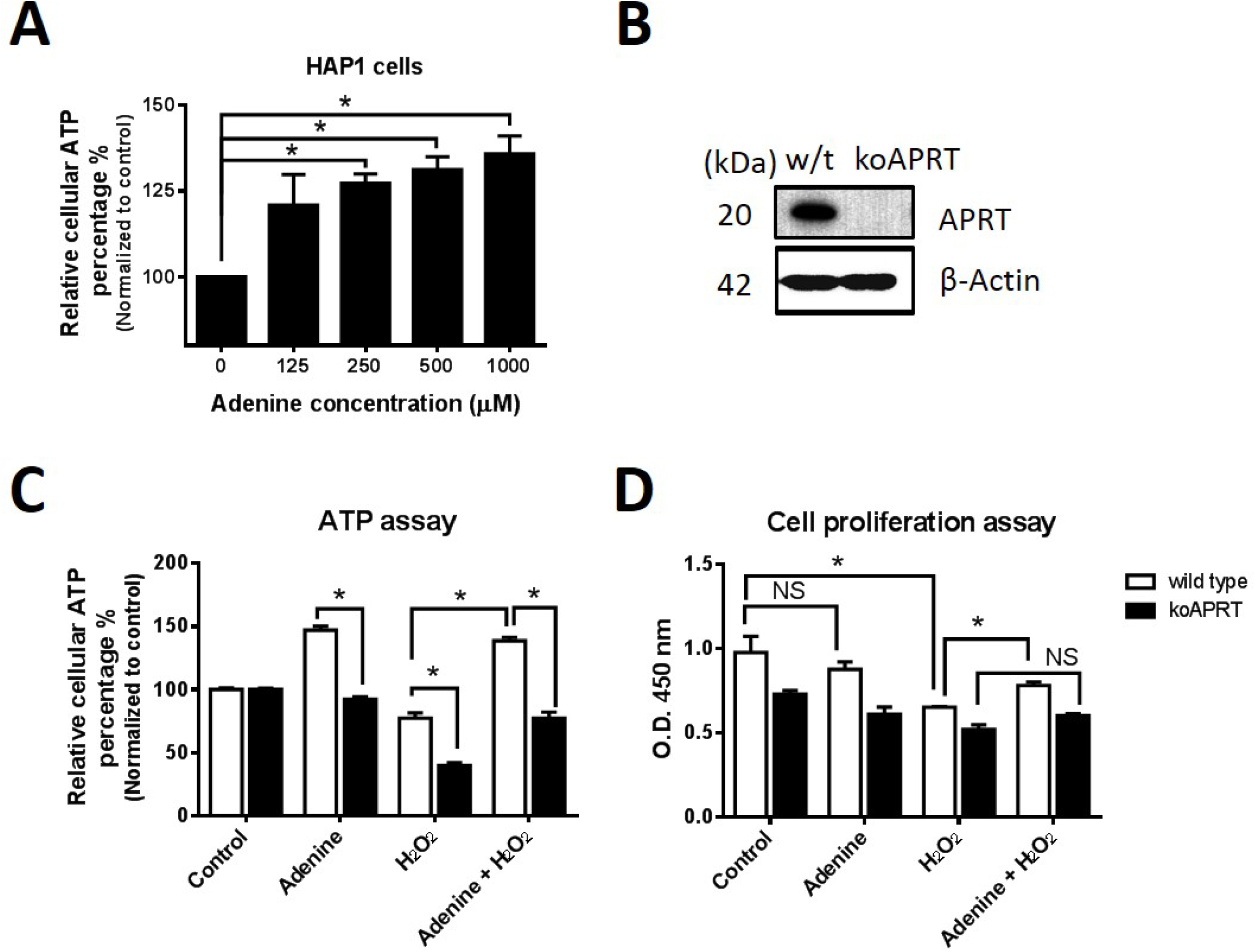
APRT mediated the elevation of cellular ATP levels to prevent hydrogen peroxide-induced oxidative damage in HAP1 cells. (A) Exposure of adenine increased the cellular ATP in a dose-dependent manner. (B) Representative Western immunoblots for APRT in wild-type and APRT knockout HAP1 cells; β-actin was used as loading control. (C) APRT mediated ATP elevation reduced the impact of ATP decline under hydrogen peroxide treatment. (D) Adenine protected the hydrogen peroxide-induced cell damage by CCK8 measurements. Data were expressed as mean ± SEM. *P < 0.05 vs. vehicle, N=5.

### Adenine elevated the ATP level and accelerated the cell migrations

Fibroblasts and keratinocytes are the main cell types that play essential roles in wound healing (Mansbridge, Liu et al., 1999). In the presence of wounds, fibroblasts migrate to the wound site, and keratinocytes proliferate to reform the epidermis to seal the wound. Here, we tested the effect of adenine on cell migration in NIH-3T3 fibroblasts and NHEK, individually. In NIH-3T3 fibroblasts, Figure 6A shows that adenine promoted ATP generation compared with the control dose dependently. The representative images from Boyden chamber assay (Jin, Xiao et al., 2015) revealed that the migratory number of adenine-treated cells also increased exponentially in a vertical orientation compared with that of the control group (135 ± 15 cells/HPF in 500 μmol adenine vs. 40 ± 5 cells/HPF in the control group, P < 0.05, Figure 6B). Similarly, adenine elevated the cellular ATP and accelerated the cellular re-epithelization in NHEK cells by a horizontal migration assay (Figure 7). The average healed area for NHEK cells compared with the initial areas at 48 h was 65.6% ± 4.5% in the adenine-treated group, whereas I the control group, it was 35.2% ± 3.6% (Figure 7B).

**Fig. 6.**
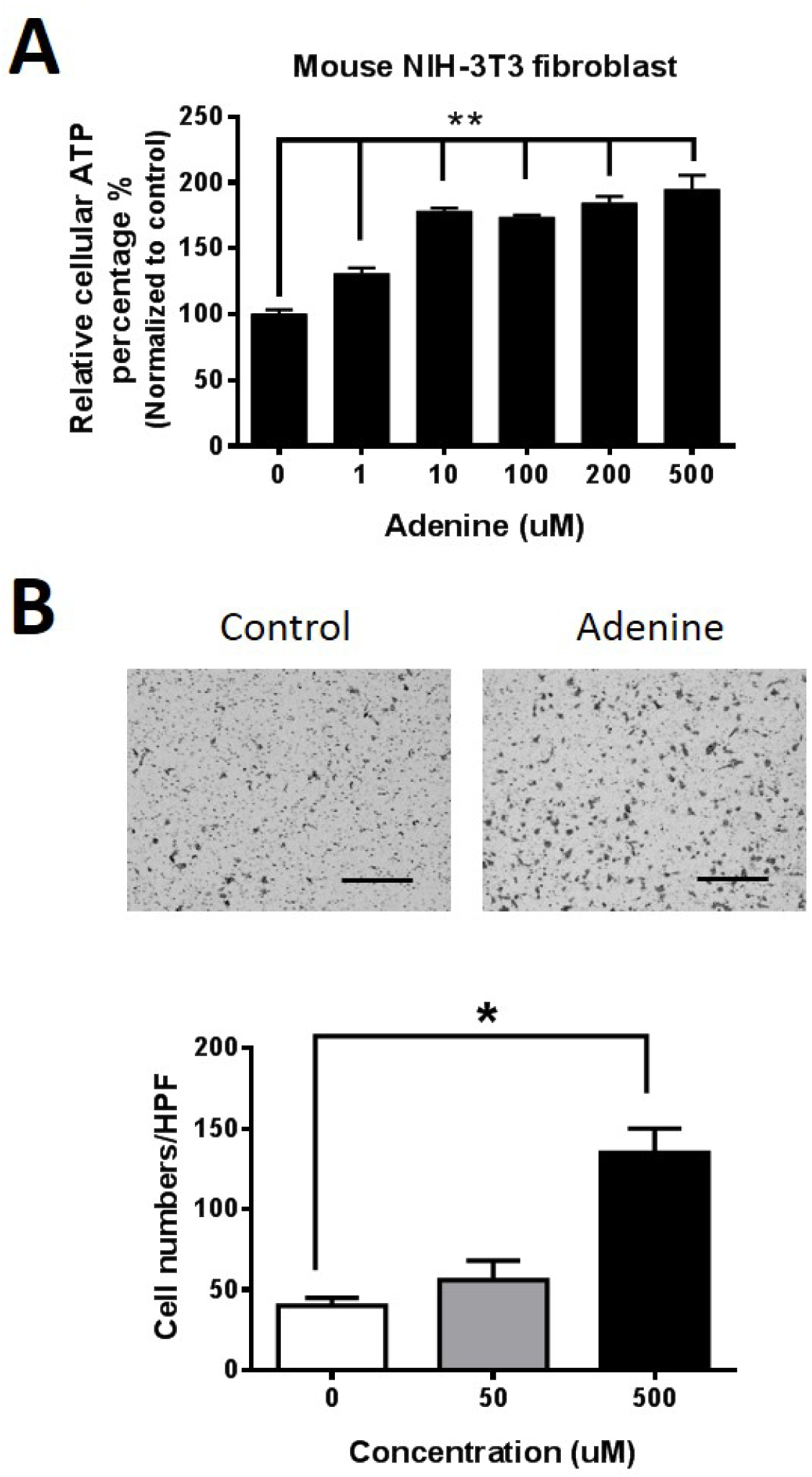
Effects of exogenous adenine on the vertical migration of NIH-3T3 fibroblasts. (A) Exposure of adenine increased the cellular ATP levels in a dose-dependent manner in mouse NIH-3T3 fibroblasts. (B) Transmigration assay by Boyden chambers for NIH-3T3 fibroblasts under adenine exposure. Data were expressed as mean ± SEM. *, P < 0.05; **, P < 0.01 vs. control group.

**Fig. 7.**
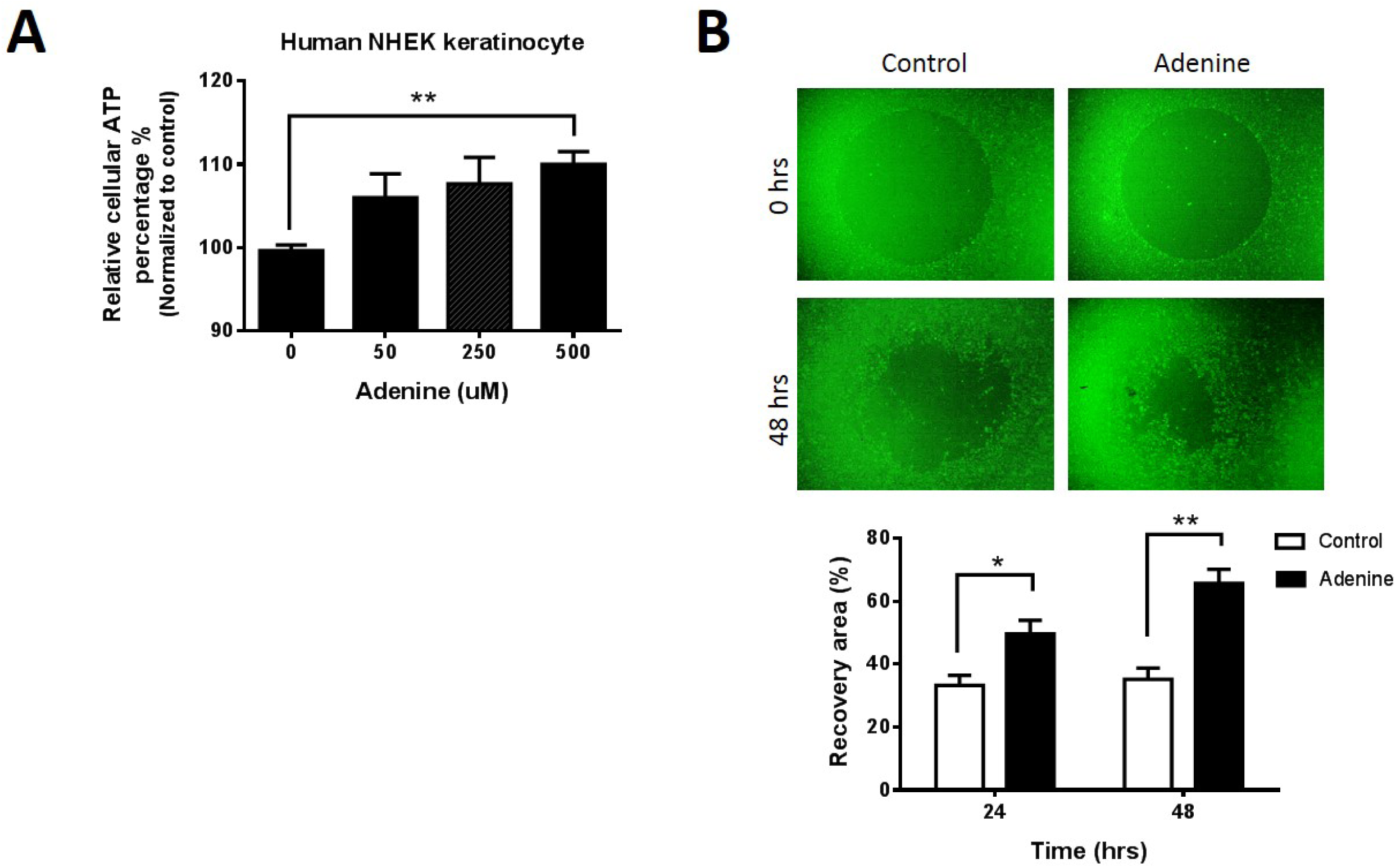
Effects of exogenous adenine on the horizontal migration of normal human epithelial keratinocytes (NHEK). **(A)** Exposure of adenine increased the cellular ATP levels in a dose-dependent manner in NHEK cells. (B) Oris cell migration assay for NHEK cells. Data were expressed as mean ± SEM. *, P < 0.05; **, P < 0.01 vs. control group.

## Discussion

Wound healing requires energy for the proliferation, migration, and remodeling of cells by fibroblasts and keratinocytes to close the wound in the epidermal layers of the skin (Wojtowicz, Oliveira et al., 2014). The proliferating epithelial cells and fibroblasts, as well as the wounded skin, primarily depend on the glycolytic pathway for energy requirement (Im & Hoopes, 1970b). However, in the wound of patients with diabetes, cellular injury and ischemia can lead to a decline in the levels of energetic metabolites, including ATP and phosphocreatine. These concentrations of energy metabolites in the wounds of patients with diabetes were significantly lower than those of normal patients (Smith, Mills et al., 1999). Given that energy is required for every aspect of the wound healing process, many strategies sustain the cellular ATP level by encapsulated ATP or blocked ATP loss would be developed as medications for morphogenetically active wound-healing formulations (Muller, Wang et al., 2017, Sarojini, Billeter et al., 2017). The cellular nucleotide level was under strict control in normal conditions (Wang, Wang et al., 2016); once a tissue sustains injury, enzymes involved in either salvage or *de novo* pathway were up-regulated for nucleotide production (Im & Hoopes, 1970a). In contrast to *de novo* nucleotide biosynthesis, the salvage pathway is extremely favorable for cells. The salvage pathway is mainly activated through growth factors, including keratinocyte growth factor and epidermal growth factor (Lanahan, Williams et al., 1992), by positively regulating the nucleotide production to facilitate wound healing (Gassmann et al., 1999). Our results revealed that APRT was up-regulated in cutaneous wound healing both in wild-type and diabetic mice. Interestingly, Western analysis from diabetic mice showed that the elevated APRT protein in the control group seemed slightly higher than that in the adenine-treated group (Figure 2F). Hence, the amplitude of APRT expression might correlate with the demands of adenylate nucleotides on the duration of wound repair. Considering that the rate of wound closure in the control group was slower than that in the adenine group (Figure 2B), the demand of APRT was higher in the control group for the accelerating wound repair.

Logically, APRT-mediated purine salvage is a process of energy consumption, including the synthesis of AMP and the interconversion of adenylate nucleotides by adenylate kinase (Figure 3)(Cheng, Young et al., 2016, Cheng, Young et al., 2015, Young & Chen, 2015, Young et al., 2015). Previous studies investigated the effect of adenine to measure the ATP solely, and rarely applied the concept of energy charge to distinguish from the changes among adenylate nucleotides. In this study, we quantified the time course of adenylate nucleotides in NIH-3T3 fibroblasts. Our result showed that the AMP increased immediately by exogenous adenine addition in 10 min, followed by ADP and then ATP (Table 1). This phenomenon exhibited the role of APRT-mediated purine salvage, especially elevated the amplitude of AMP and reduced the AEC, indicating that the energy status of cells was low by adenine addition. However, this phenomenon is contradictory to extensive experiment studies, in which adenine elevated the cellular ATP levels both *in vivo* and *in vitro*. In this study, our data explained that exogenous adenine could enlarge the cellular adenylate nucleotide pool (AMP + ADP + ATP). Despite the increasing amplitude of ATP is less than AMP, the total ATP after adenine treatment was also significantly increase compare with control. Furthermore, the amplitude of increase AMP made the decrease of AEC, which represent the low energy status in the cell. The phenomenon may triggered the catabolic ATP-producing pathways, such AMPK activation to promote glucose sparing and oxidative metabolism (Supplemental Figure 2).

The ATP molecule released from an injured tissue may act as a signal to affect cellular responses, thereby accelerate wound healing (Yin, Xu et al., 2007). However, the extracellular ATP that triggered purinergic receptor signaling is differing from adenine, even though both of them mediated through an ATP molecule. The ATP molecule carries four negative charges; thus, exogenous ATP cannot pass through the cell membrane directly. As for adenine, which is water soluble and can transport across the cell membranes or bind to highly specific receptor proteins into the cytoplasm (Cartier, 1980). Adenine, as one of the main substrates used for storing of RBCs in a blood bag (Mazor, Dvilansky et al., 1994, Paglia, Sigurjonsson et al., 2016), and the synthesis of nucleotides in RBC occurs only through the salvage pathway (Baranowska-Bosiacka, Dziedziejko et al., 2009). Our study has utilized the characteristic of APRT to convert adenine into AMP and enlarge the cellular adenylate nucleotide pool.

Previously, we demonstrated a link between adenine and AMPK in many cell models, and the underlying mechanism may be via APRT anabolism to increase the cellular AMP level to active AMPK, subsequently enhancing ATP generation (Cheng et al., 2015, Young & Chen, 2015, Young et al., 2015). In fact, topical adenine could also slightly elevate the phosphorylation of AMPK in the wound of diabetic mice compare with vehicle group (Supplementary Figure 2). Those data provided an evidence *in vivo* that adenine may accelerate diabetic wound healing through the AMPK signaling pathway.

On the other hand, ROS arousing to oxidative stress is a critical issue in the pathogenesis of chronic wounds. The effect of hydrogen peroxide in mammalian cells accompanied by a marked reduction of the ATP level and AEC through the glyceraldehyde-3-phosphate dehydrogenase inhibition (Lelli, Becks et al., 1998, Stuart & Holmsen, 1977). Our data showed that ATP level reduction by hydrogen peroxide reversed by adenine and markedly prolonged cell viability. The underlying mechanism is via the APRT-mediated purine salvage to reduce the impairment of hydrogen peroxide in ATP levels and cell viability.

Taken together, our data indicated the role of APRT during diabetic wound healing by regulating the nucleotide pool after tissue injury and demonstrated the improvement by topical adenine, which highlights its value as a promising agent in therapeutic intervention. This study provides an explanation for the up-regulation of APRT in tissues and adenine supplementation that resulted in an enlargement of the adenylate nucleotides for ATP generation.

## Materials and Methods

### Reagents

Adenine was purchased from Sigma-Aldrich (St. Louis, MO, USA). Dulbecco’s modified Eagle’s medium (DMEM), trypsin, penicillin-streptomycin, and Newborn calf serum (NBCS) were from Invitrogen™ (Carlsbad, CA, USA).

*Cell culture.* Mouse NIH-3T3 fibroblasts were prepared as described previously (Young et al., 2015). Cells were maintained in culture flasks containing growth medium (DMEM with 1.5 g/L sodium bicarbonate, 100 U/mL penicillin-streptomycin and 10% FBS) at 37°C in a 100% humidified atmosphere with 5% CO2 (Young et al., 2015). HAP1 knockout cell lines were generated by Horizon Discovery (Cambridge, UK) (Essletzbichler, Konopka et al., 2014).

### Preparation of working standard solutions

To study the effect of exogenous adenine in cellular adenylate balance, 1 × 10^5^ NIH-3T3 fibroblasts exposed to medium containing different concentrations of adenine were cultured in 24 well plates for the indicated time. After removal of the supernatant medium, 1 mL of 50% ACN containing 50 ppb fludarabine phosphate was used as an internal standard (IS) to add to the cells. The cell lysate was transferred to a microcentrifuge tube and was centrifuged at 12,000 rpm for 10 minutes at 4°C to remove the pellet. The supernatant was collected, lyophilized, and stored at −80°C until used.

### Instrumental and analytical conditions

The analysis of adenine and its derivative metabolites was achieved by an Agilent HPLC 1200 series system (Agilent, USA) coupled with an API 4000 triple quadrupole mass spectrometer (Applied Biosystems, MDS Sciex, Canada). The API 4000 triple quadrupole mass spectrometer is equipped with a Turbo V ion source with electrospray ionization (ESI) and was operated in negative ion mode. Multiple reactions monitoring (MRM) mode was applied for the quantification of analytes. The compound dependent parameters for each MRM transition of analytes are given in Supplement Table.1 and the data was collected using the Analyst 1.6.2 Software (AB Sciex).

### Measurement of intracellular ATP

The cellular ATP levels were quantified in 100 μL ATP assay buffer or from 20-30 mg wound skin harvested in 200 μL assay buffer and lysed with the bead shocker following the manufacturer’s instructions (CellTiter-Glo luminescent ATP assay kit, Promega, Madison). The protein content from identical treated cells or tissues were determined by BCA Protein Assay kit for normalization (Young, Huang et al., 2014).

### Vertical migration of NIH-3T3 fibroblasts

Cell migration was assessed using a modified Boyden chamber (8 μm pores; Transwell, Corning, Massachusetts)(Wu, Young et al., 2013). NIH-3T3 fibroblasts in 100 μL serum-free DMEM were plated into the upper chamber at 2.5 × 10^5^ cells/well, while the lower champer contained 30 μL DMEM including 10% FBS. Chambers were incubated at 37°C for 18 hours. After incubation, the migrated cells on the lower surface of the membrane were fixed, stained, and quantified with the image J software.

### Horizontal migration of normal human epithelial keratinocytes (NHEK)

The migration of NHEKs were real time monitored using Oris(tm) cell migration assay system (Gough, Hulkower et al., 2011). NHEKs labelled with CMFDA were plated in the wells of a 96- well Oris(tm) plate. After a 24 hour attachment period, a void was created by removing stoppers from all test wells. For pre-migration controls, stoppers remained in wells until assay readout. Cells migration were quantified by measurement of the void area by digital camera.

### Animal and wound model

All experimental procedures were approved by the Institutional Animal Care and use Committee of the University of Fu-Jen Catholic University. Eight-week-old male wild-type (C57BL/6) and diabetic (BKS Cg-Dock7m^+/+^ Leprdb/JNar1) mice were purchased from the National Laboratory Animal Center (Taipei, Taiwan). Mice was anaesthetized (i.p. injection) using a combination of Zoletil 50®, Rompun® and saline (a ratio of 1:1:2, respectively). Before surgery, the hair was shaved from the dorsal surface of mice. Single full-thickness excisional skin wound was created on mice by using an 8 mm diameter biopsy punch, and the excised skin tissues were used as control. A donut-shape silicone splint was stitched around the wound with an immediate-bonding adhesive (Dunn, Prosser et al., 2013) and the wound areas were topically applied with 20 μL hydrogel daily. After drug administration, a semi-occlusive dressing (Tegaderm, 3M, St. Paul, MN, USA) was applied to cover the wound and splint. Images of wound were acquired daily with a digital camera. Time to wound closure was defined as the time needed for the wound bed to be completely re-epithelialized. At the day that mice were sacrificed, the wound skin samples were harvested using a 8-mm biopsy punch.

### Histological analysis

Mouse skin tissues containing wound area were fixed in 10% formaldehyde for paraffin embedding. The sections of paraffin-embedded tissue were stained with Masson’s trichrome stain kit (ScyTek Laboratories). For immunohistochemical staining, a non-biotin-amplified kit (Novolink, Novocastra Laboratories) was used according to the manufacturer’s instruction (Young et al., 2014). Sections were washed and incubated for 1 hour in the presence of appropriate secondary antibodies. Image analysis was obtained using a Nikon digital system.

### Cell viability assay

Cell viability was determined by the cell counting kit-8 (CCK-8) (Dojindo Molecular Technologies, Kumamoto, Japan). Briefly, 10,000 cells were plated in 96-well plates overnight. After drug treatment, CCK-8 reagent added into the plate and incubated in CO2 incubator at 37°C for additional 2 hours (Young, Tang et al., 2019). The absorbance was measured at 450 nm using a microplate reader.

### Protein extraction and western blot

Cell lysates were prepared as described previously (Young et al., 2014). In brief, cells were dissolved in RIPA buffer containing a protease inhibitor cocktail (Roche Applied Science, Indianapolis, IS, USA) were quantitated using the BCA protein assay kit (Energenesis- Biomedical, Taipei, TW)(Young, Chen et al., 2006). Mouse skin tissue was homogenized in T-PER Tissue Protein Extraction Reagent (Thermo Fisher Scientific) plus a protease inhibitor cocktail (Roche Applied Science, Indianapolis, IS, USA) using a cooled bead shocker SpeedMill (AnalytikJena, Jena, Germany). Equal amount of protein was separated by SDS-PAGE, and transfer to PVDF membranes (Millipore, Billerica, MA, USA). Membranes were blocked with 5% bovine serum albumin (BSA) in PBS and incubated with anti-APRT antibody or an anti-β- Actin antibody (Cell Signaling Technology, USA) at 4°C overnight followed by the corresponding secondary antibody for 1 hr at room temperature. Immunoreactive bands were incubated with LumiFlash Prime Chemiluminescent substrate and recorded with a Chemlux SPX-600 Series Imaging Systems (Energenesis-Biomedical, Taipei, TW)(Sun, Young et al., 2015). The detected signals were quantified using ImageJ software (http://image.nih.gov/ij/).

### Statistical analysis

Experimental results are expressed as the mean±SEM, and comparisons between the data sets were performed by non-parametric Mann Whitney U or Kruskall Wallis tests (SPSS & GraphPad Prism 6; GraphPad Software, La Jolla, CA). Statistical difference was considered to be significant when P<0.05.

## Supplementary Materials

Fig. S1. The expression of APRT during cutaneous wound repair in wild-type mice.

Fig. S2. The change of AMPK phosphorylation during diabetic wound healing

## Acknowledgments

We are thankful for the support of the Spectral Confocal Laser Scanning Platform Laboratory in department of Life Science, Fu-Jen Catholic University.

## Author contributions

G.H. Young, J.T. Lin. and H.M. Chen. conceived the study. C.F. Ho, Q.Y. Kuok, R.C. Hsu, W.R. Liao, and C.C. Chen performed the experiments. Y.F. Cheng analyzed the data. G.H. Young wrote the paper. J.T. Lin. and H.M. Chen. supervised the study.

## Data and materials availability

The HAP1 of APRT knockout cells are available from H.M. Chen. under a material transfer agreement with the company of Energenesis-Biomedical.

## Figures

Table 1 **Time course of the adenylate nucleotide response to adenine supplementation in NIH-3T3 fibroblasts.** Absolute concentrations of adenylate nucleotides exposed to different amounts of adenine, as monitored by MRM methods; N = 4. Data were presented as mean ± SEM. *, P < 0.05; ^&^, P < 0.05; ^$^, P < 0.05, significantly different from 0 μM adenine control in each time group.

## Supplementary Materials

**Fig. S1. The expression of APRT during cutaneous wound repair in wild-type mice.** (A) The representative images of wound healing in mice at indicated time. (B) Representative western immunoblots for APRT in C57BL/6 mice skin at indicated time point, β-actin were used as loading controls. (C) Quantitation of the intensity of western blots for APRT. Data were expressed as mean±SEM, N=3. *, P<0.05; N.S., non-significant.

**Fig. S2. The change of AMPK phosphorylation during diabetic wound healing.** (A) Representative western immunoblots for P-AMPK and T-AMPK treated with adenine in diabetic wounds at day 20 post-injury. β-actin were used as loading controls. (B) Quantitation of the intensity of western blots for P-AMPK. Data were expressed as mean±SEM. *, P<0.0. (C) Histological sections were stained by immunohistochemistry with anti-P-AMPK antibodies and were counterstained with Haematoxylin. Scale bars, 500 & 50 μm, respectively. 5 versus control skin; **, P<0.01 versus control skin, N=5.

## References

Andreea SI, Marieta C, Anca D (2008) AGEs and glucose levels modulate type I and III procollagen mRNA synthesis in dermal fibroblasts cells culture. Exp Diabetes Res 2008: 473603

Atkinson DE (1968) The energy charge of the adenylate pool as a regulatory parameter. Interaction with feedback modifiers. Biochemistry 7: 4030–4

Baranowska-Bosiacka I, Dziedziejko V, Safranow K, Gutowska I, Marchlewicz M, Dolegowska B, Rac ME, Wiszniewska B, Chlubek D (2009) Inhibition of erythrocyte phosphoribosyltransferases (APRT and HPRT) by Pb2+: a potential mechanism of lead toxicity. Toxicology 259: 77–83

Bhagavan NV, Ha C-E (2011) Essentials of medical biochemistry: with clinical cases. Elsevier/Academic Press, Amsterdam; Boston

Cano Sanchez M, Lancel S, Boulanger E, Neviere R (2018) Targeting Oxidative Stress and Mitochondrial Dysfunction in the Treatment of Impaired Wound Healing: A Systematic Review. Antioxidants (Basel) 7

Carette JE, Raaben M, Wong AC, Herbert AS, Obernosterer G, Mulherkar N, Kuehne AI, Kranzusch PJ, Griffin AM, Ruthel G, Dal Cin P, Dye JM, Whelan SP, Chandran K, Brummelkamp TR (2011) Ebola virus entry requires the cholesterol transporter Niemann-Pick C1. Nature 477: 340–3

Cartier PH (1980) Adenine uptake by isolated rat thymocytes. J Biol Chem 255: 4574–82

Cheng YF, Young GH, Chiu TM, Lin JT, Huang PR, Kuo CY, Liang YJ, Chen PK, Chen SF, Cheng CY, Chen HM (2016) Adenine supplement delays senescence in cultured human follicle dermal papilla cells. Exp Dermatol 25: 162–4

Cheng YF, Young GH, Lin JT, Jang HH, Chen CC, Nong JY, Chen PK, Kuo CY, Kao SH, Liang YJ, Chen HM (2015) Activation of AMP-Activated Protein Kinase by Adenine Alleviates TNF-Alpha-Induced Inflammation in Human Umbilical Vein Endothelial Cells. PLoS One 10: e0142283

Dunn L, Prosser HC, Tan JT, Vanags LZ, Ng MK, Bursill CA (2013) Murine model of wound healing. J Vis Exp: e50265

Essletzbichler P, Konopka T, Santoro F, Chen D, Gapp BV, Kralovics R, Brummelkamp TR, Nijman SM, Burckstummer T (2014) Megabase-scale deletion using CRISPR/Cas9 to generate a fully haploid human cell line. Genome Res 24: 2059–65

Falanga V (2005) Wound healing and its impairment in the diabetic foot. Lancet 366: 1736–43

Foker JE (1988) Method for stimulating recovery from ischemia. In Google Patents

Frenguelli BG (2017) The Purine Salvage Pathway and the Restoration of Cerebral ATP: Implications for Brain Slice Physiology and Brain Injury. Neurochem Res

Gassmann MG, Stanzel A, Werner S (1999) Growth factor-regulated expression of enzymes involved in nucleotide biosynthesis: a novel mechanism of growth factor action. Oncogene 18: 6667–76

Gough W, Hulkower KI, Lynch R, McGlynn P, Uhlik M, Yan L, Lee JA (2011) A quantitative, facile, and high-throughput image-based cell migration method is a robust alternative to the scratch assay. J Biomol Screen 16: 155–63

Hardie DG (2004) The AMP-activated protein kinase pathway--new players upstream and downstream. J Cell Sci 117: 5479–87

Hardie DG (2015) AMPK: positive and negative regulation, and its role in whole-body energy homeostasis. Curr Opin Cell Biol 33: 1–7

Howard JD, Sarojini H, Wan R, Chien S (2014) Rapid granulation tissue regeneration by intracellular ATP delivery--a comparison with Regranex. PLoS One 9: e91787

Im MJ, Hoopes JE (1970a) Energy metabolism in healing skin wounds. J Surg Res 10: 459–64

Im MJ, Hoopes JE (1970b) Enzyme activities in the repairing epithelium during wound healing. J Surg Res 10: 173–9

Jin CE, Xiao L, Ge ZH, Zhan XB, Zhou HX (2015) Role of adiponectin in adipose tissue wound healing. Genet Mol Res 14: 8883–91

Kamatani N, Carson DA (1981) Dependence of adenine production upon polyamine synthesis in cultured human lymphoblasts. Biochim Biophys Acta 675: 344–50

Lanahan A, Williams JB, Sanders LK, Nathans D (1992) Growth factor-induced delayed early response genes. Mol Cell Biol 12: 3919–29

Lelli JL, Jr., Becks LL, Dabrowska MI, Hinshaw DB (1998) ATP converts necrosis to apoptosis in oxidant-injured endothelial cells. Free Radic Biol Med 25: 694–702

Leu JG, Chiang MH, Chen CY, Lin JT, Chen HM, Chen YL, Liang YJ (2017) Adenine accelerated the diabetic wound healing by PPAR delta and angiogenic regulation. Eur J Pharmacol 818: 569–577

Lin JT, Chen HM, Chiu CH, Liang YJ (2014) AMP-activated protein kinase activators in diabetic ulcers: from animal studies to Phase II drugs under investigation. Expert Opin Investig Drugs 23: 1253–65

Mansbridge JN, Liu K, Pinney RE, Patch R, Ratcliffe A, Naughton GK (1999) Growth factors secreted by fibroblasts: role in healing diabetic foot ulcers. Diabetes Obes Metab 1: 265–79

Mazor D, Dvilansky A, Meyerstein N (1994) Prolonged storage of red cells: the effect of pH, adenine and phosphate. Vox Sang 66: 264–9

Muller WEG, Wang S, Wiens M, Neufurth M, Ackermann M, Relkovic D, Kokkinopoulou M, Feng Q, Schroder HC, Wang X (2017) Uptake of polyphosphate microparticles in vitro (SaOS-2 and HUVEC cells) followed by an increase of the intracellular ATP pool size. PLoS One 12: e0188977

Murray AW (1971) The biological significance of purine salvage. Annu Rev Biochem 40: 811–26

Nouvong A, Ambrus AM, Zhang ER, Hultman L, Coller HA (2016) Reactive oxygen species and bacterial biofilms in diabetic wound healing. Physiol Genomics 48: 889–896

Paglia G, Sigurjonsson OE, Bordbar A, Rolfsson O, Magnusdottir M, Palsson S, Wichuk K, Gudmundsson S, Palsson BO (2016) Metabolic fate of adenine in red blood cells during storage in SAGM solution. Transfusion 56: 2538–2547

Papanas N, Maltezos E (2010) Benefit-risk assessment of becaplermin in the treatment of diabetic foot ulcers. Drug Saf 33: 455–61

Pastar I, Stojadinovic O, Yin NC, Ramirez H, Nusbaum AG, Sawaya A, Patel SB, Khalid L, Isseroff RR, Tomic-Canic M (2014) Epithelialization in Wound Healing: A Comprehensive Review. Adv Wound Care (New Rochelle) 3: 445–464

Rossomando EF, Bertolami CN (1983) Alterations in purine salvage and hypoxanthine levels in granulation tissue during skin wound repair. J Surg Res 35: 259–63

Sarojini H, Billeter AT, Eichenberger S, Druen D, Barnett R, Gardner SA, Galbraith NJ, Polk HC, Jr., Chien S (2017) Rapid tissue regeneration induced by intracellular ATP delivery-A preliminary mechanistic study. PLoS One 12: e0174899

Smith DG, Mills WJ, Steen RG, Williams D (1999) Levels of high energy phosphate in the dorsal skin of the foot in normal and diabetic adults: the role of 31P magnetic resonance spectroscopy and direct quantification with high pressure liquid chromatography. Foot Ankle Int 20: 258–62

Stuart MJ, Holmsen H (1977) Hydrogen peroxide, an inhibitor of platelet function: effect on adenine nucleotide metabolism, and the release reaction. Am J Hematol 2: 53–63

Sun CY, Young GH, Hsieh YT, Chen YH, Wu MS, Wu VC, Lee JH, Lee CC (2015) Protein-bound uremic toxins induce tissue remodeling by targeting the EGF receptor. J Am Soc Nephrol 26: 281–90

Thimm D, Schiedel AC, Peti-Peterdi J, Kishore BK, Muller CE (2015) The nucleobase adenine as a signalling molecule in the kidney. Acta Physiol (Oxf) 213: 808–18

Traut TW (1994) Physiological concentrations of purines and pyrimidines. Mol Cell Biochem 140: 1–22

Vande Berg JS, Robson MC (2003) Arresting cell cycles and the effect on wound healing. Surg Clin North Am 83: 509–20

Wang J, Wan R, Mo Y, Li M, Zhang Q, Chien S (2010) Intracellular delivery of adenosine triphosphate enhanced healing process in full-thickness skin wounds in diabetic rabbits. Am J Surg 199: 823–32

Wang X, Wang G, Li X, Fu J, Chen T, Wang Z, Zhao X (2016) Directed evolution of adenylosuccinate synthetase from Bacillus subtilis and its application in metabolic engineering. J Biotechnol 231: 115–121

Wojtowicz AM, Oliveira S, Carlson MW, Zawadzka A, Rousseau CF, Baksh D (2014) The importance of both fibroblasts and keratinocytes in a bilayered living cellular construct used in wound healing. Wound Repair Regen 22: 246–55

Wu VC, Young GH, Huang PH, Lo SC, Wang KC, Sun CY, Liang CJ, Huang TM, Chen JH, Chang FC, Chen YL, Kuo YS, Chen JB, Chen JW, Chen YM, Ko WJ, Wu KD, group N (2013) In acute kidney injury, indoxyl sulfate impairs human endothelial progenitor cells: modulation by statin. Angiogenesis 16: 609–24

Yin J, Xu K, Zhang J, Kumar A, Yu FS (2007) Wound-induced ATP release and EGF receptor activation in epithelial cells. J Cell Sci 120: 815–25

Young G-H, Chen H-M (2015) Adenine Supplement Suppresses Diabetic Nephropathy Through AMPK and Sirt-1 Pathways. Hong Kong Journal of Nephrology 2: S19–S20

Young GH, Chen HM, Lin CT, Tseng KC, Wu JS, Juang RH (2006) Site-specific phosphorylation of L-form starch phosphorylase by the protein kinase activity from sweet potato roots. Planta 223: 468–78

Young GH, Huang TM, Wu CH, Lai CF, Hou CC, Peng KY, Liang CJ, Lin SL, Chang SC, Tsai PR, Wu KD, Wu VC, Ko WJ, group N (2014) Hemojuvelin modulates iron stress during acute kidney injury: improved by furin inhibitor. Antioxid Redox Signal 20: 1181–94

Young GH, Lin JT, Cheng YF, Huang CF, Chao CY, Nong JY, Chen PK, Chen HM (2015) Identification of adenine modulating AMPK activation in NIH/3T3 cells by proteomic approach. J Proteomics 120: 204–14

Young GH, Tang SC, Wu VC, Wang KC, Nong JY, Huang PY, Hu CJ, Chiou HY, Jeng JS, Hsu CY (2019) The functional role of hemojuvelin in acute ischemic stroke. J Cereb Blood Flow Metab: 271678×19861448

